# Modeling tissue growth with the Stokes equation

**DOI:** 10.1101/641282

**Authors:** Teemu J. Häkkinen, Jukka Jernvall, Antti Hannukainen

## Abstract

We present a cell-free continuum model for simulating generalized bulk tissue growth in 3D. We assume that the tissue behaves mechanically as viscous fluid so that its behavior can be described with the Stokes equation with mass sources. The growth is directed by a diffusing morphogen produced by specialized signaling centers, whose positions are established through a reaction-diffusion system coupled with differentiation. We further assume that the tissue interface may be stiff (modeled as surface tension), and that tissue adhesion can vary (modeled as variable viscosity). The numerical validity of the implementation is investigated using test cases with known solutions, and the model dynamics are demonstrated in simulations of idealized tissue growth. The combination of Stokes equation and diffusing morphogens allow the integration of patterning and growth as in real organs systems such as limbs and teeth. We propose that the presented techniques could be useful for simulating and exploring mechanistic principles of tissue growth in various developing organs.

## 1 Introduction

One of the most challenging questions in computational biology is how to simulate tissue morphogenesis, or the formation of organized, non-homogenous tissues giving rise to organs. At the highest level of abstraction the existing modeling approaches can be divided into two categories: Cell-based and continuum models [1, 2]. In cell-based models the computational nodes are more or less explicitly identified as cells, whereby cell-specific properties such as cell shape, adhesion and cell-cell signaling are the main targets of modeling. The cell-based approach could be characterized as bottom-up, where the macro (or tissue-level) properties are obtained as a result of simulating a large number of micro (or cell-level) phenomena.

Continuum models take the opposite view: The objective is to simulate the behavior and the properties of the tissue directly by making a continuum assumption, that is, assuming that macro-level behavior of a large number of cells can be described directly, ignoring the cell-level phenomena. The continuum approach by definition imposes a lower-bound limit on the level of detail that can be represented with such models. On the other hand, continuum models can usually be expected to have fewer free parameters, and also have significant scaling advantages when attempting to simulate large systems consisting of thousands of cells.

In this paper we investigate the application of the continuum approach in modeling generalized bulk tissue growth in 3D using the Stokes equation. Our model draws inspiration from previous Stokes models by Murea et. al. [3] and Boehm et. al. [4] that considered prescribed growth patterns in 2D and 3D, respectively. We augment these previous models by including of a dynamic patterning mechanism with differentiation for establishing the tissue growth centers, and using a form of the Stokes equation with variable viscosity. The differentiation mechanism, similar in principle to what was implemented in [5], mimics the state changes that growing tissues undergo (e.g., signaling centers in teeth [6]). Variable viscosity models the changes in the adhesive properties of the tissue, such as changes in cell adhesion, which has been proposed to be an important factor in morphogenesis [7]. We also represent the tissue-boundary as an implicit surface using the level set method [8], rather than explicitly through mesh deformations, which simplifies the model implementation and reduces computational demands.

The paper is organized as follows: We first describe the techniques and the main components of the model, including patterning, tissue growth dynamics and interface tracking. We then demonstrate the numerical validity of the implementation through simulations in cases where the correct solution is known. Finally, we demonstrate the virtual tissue growth in a few test cases and briefly discuss potential applications.

## 2 Methods

We consider the growth of a bulk tissue contained within a static computational domain Ω. The parts of Ω occupied by the tissue are denoted by Ω_2_(*t*) (the interior), with the rest of Ω denoted by Ω_1_(*t*) (the exterior). In practice, the exterior will often represent a tissue as well, but in this paper we restrict our focus on a case where all active growth takes place within the interior, and the exterior serves the role of a passive surrounding material only. The initial shape of Ω_2_(*t*) within Ω can be chosen as needed, for example as a small sphere, with Ω typically set as the unit cube. For notational clarity, we will drop the explicit mention of time *t* in the following.

The subdomains Ω_1_ and Ω_2_ are separated by an interface Γ. Following the level set method [8], the position of Γ at time *t* is represented by the zero of a signed distance function

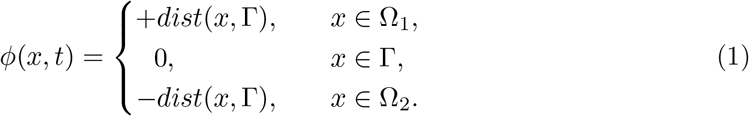

The basic outline of the model is depicted in Fig. 1. In short, we assume that a patterning mechanism acts within the tissue Ω_2_, establishing the positions of specialized signaling centers. The signaling centers produce a diffusing growth factor that both induces tissue growth and blocks further patterning locally. The growth factor determines the strength of mass sources in the Stokes equation, and the velocity field obtained by solving the resulting Stokes system is used to update the distance function (1), for the purpose of updating the position of the interface Γ. For both visualization and interface-dependent force calculation purposes, we reconstruct the implicit surface from Γ whenever the distance function (1) is updated. An algorithmic summary of the steps to solve the model is presented in Algorithm 1, with the main techniques of each step described in the following sections.

**Fig. 1.**
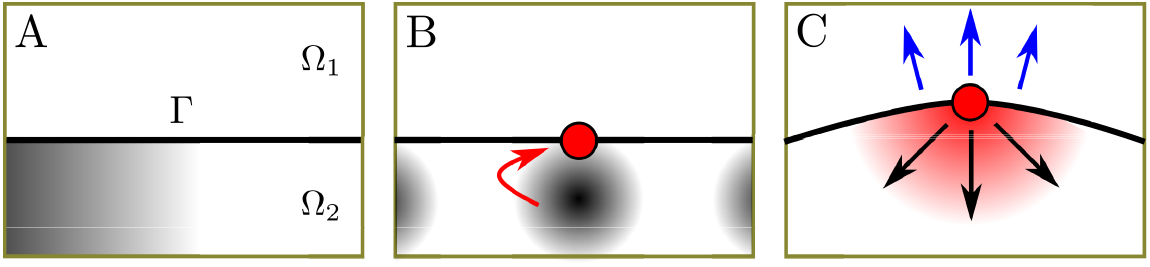
Patterning and differentiation. (A) The computational domain Ω is divided into non-overlapping subdomains of Ω_1_ (exterior) and Ω_2_ (interior), separated by interface Γ. Initially, Ω_2_ has a weak non-homogeneous concentration of the activator morphogen to start a patterning process. (B) The patterning process induces signaling centers (red) at the interface. (C) Signaling centers secrete diffusing growth factor into the interior Ω_2_, inducing growth and locally inhibiting further patterning.

We make a distinction between the ‘local time’ of each step, denoted by *t*, and the ‘global time’ of the simulation, denoted by *t*_*n*_. In practice, the global time is anchored to the growth time (Step 5 in Algorithm 1) such that at time *t*_*n*_ growth has advanced *nδ* time, where *n* is the number of iterations of Algorithm 1 and *δ* the time step. On the other hand, we assume that patterning (Step 2) happens fast relative to growth and, consequently, it is solved for longer time scales during each iteration, independent of the global simulation time.

The model equations are solved with a custom finite element method (FEM) implementation written in C++. See [10] for an introduction to FEM. The implementation is available in GitHub at https://github.com/tjhakkin/fluidtissue.

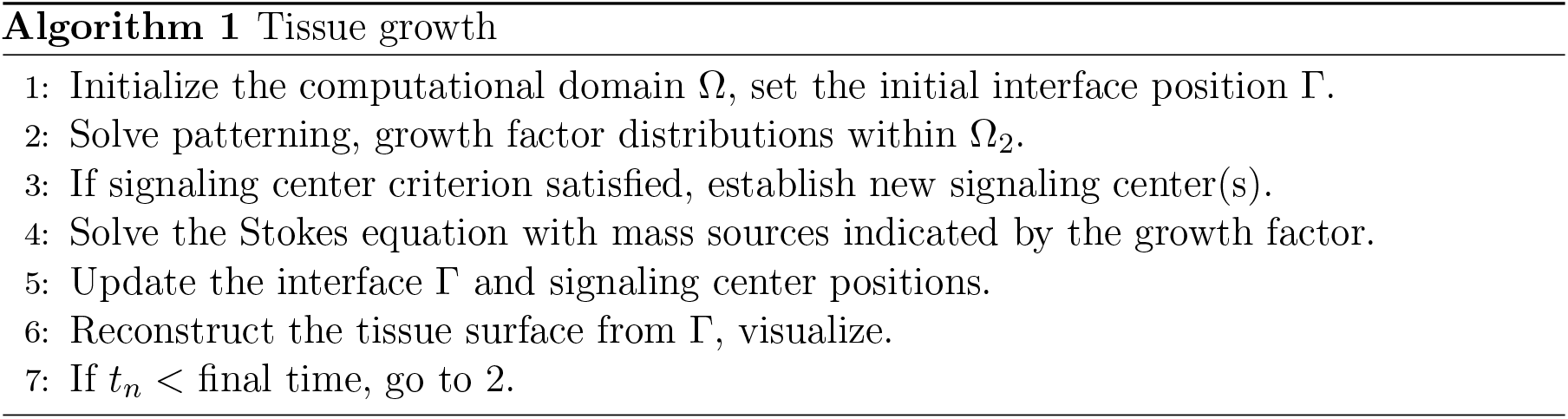

### 2.1 Patterning and growth factor

To establish the growth factor distribution within tissue Ω_2_, we use a three-morphogen reaction-diffusion system with differentiation. Two of the morphogens form a traditional activator-inhibitor loop for patterning, which induces the differentiation of signaling centers. Signaling centers produce the third morphogen, the growth factor, that both induces growth and blocks the activity of the other two morphogens. An overview of the process is shown in Fig. 1. At the initial global time *t*_*n*_ = 0, the system has no signaling centers and only weak non-homogeneous background concentration of the activator morphogen within Ω_2_. The reaction-diffusion system produces a periodic pattern of activator peaks (patterning), and upon locally reaching a threshold level of activator concentration near the interface, a signaling center is established (differentiation).

While the patterning takes place within time-dependent domain Ω_2_, the domain remains static during each patterning step. Because of this, we don’t consider morphogen advection here, but instead discuss it later when the shape of Ω_2_ is updated due to growth. We denote the concentrations of the three morphogens by *c*_*i*_(*x*, *t*) : Ω_2_×[0, *T*] → ℝ for *i* = {1, 2, 3}, with *c*_1_ the activator, *c*_2_ the inhibitor, *c*_3_ the growth factor and *T* a suitable time endpoint. Note that *T* refers to the local time of the patterning step. The general form of the governing equations for the morphogens is

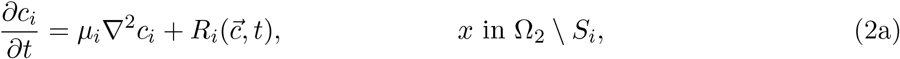

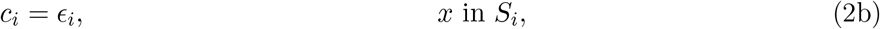

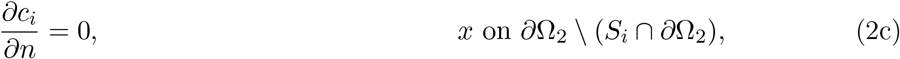

where *R*_*i*_ is the reaction term and *S*_*i*_ ⊂ Ω_2_ indicate the signaling centers producing morphogen *i* at a constant rate of *ϵ*_*i*_ ∈ ℝ^+^. In our case *S*_1_ = *S*_2_ = ∅ for all *t*. For demonstration purposes, we use the following reaction terms, extended from the classical Gierer-Meinhardt system [9]:

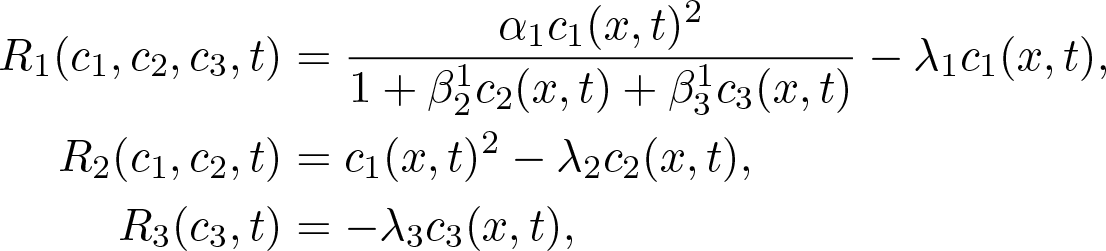

where *α*_*i*_ is the strength of morphogen *i* auto-activation, 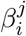 the strength of the inhibition of morphogen *i* over morphogen *j*, and *λ*_*i*_ the morphogen decay rates. All the parameters are positive scalars. With appropriate choice of initial conditions and parameters, this system is capable of producing periodic patterns, a property which we exploit here. The reaction terms used here are just an example, and in modeling real biological systems they would be chosen according to the specifics of the target system.

As is customary in FEM, we consider solving the equations in their weak forms: For *i* = {1, 2, 3}, find *c*_*i*_ in *U* (the solution space), such that

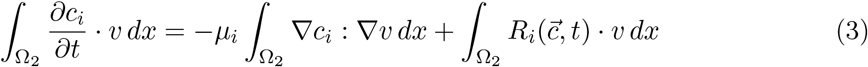

for all test functions *v* in *U*. The Neumann condition (2c) is the natural boundary condition for this problem, hence there is no need to impose any boundary conditions for *c*_1_, *c*_2_, whereas for *c*_3_ the condition (2b) needs to be included in the solution space. Note that by ‘boundary’ we refer communally to both the actual boundary and also the potential non-boundary parts of *S*_*i*_.

Denoting the standard mass and stiffness matrices by **M** and **A**, respectively, the discrete version of problem (3) can be expressed as

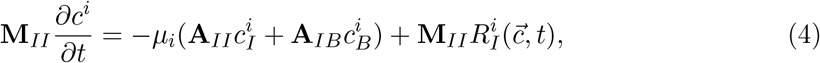

where *I* is the set of nodes corresponding to Ω_2_ \ *S*_*i*_, *B* the nodes corresponding to Ω_2_ ∩ *S*_*i*_ and *c*^*i*^ denotes the time-dependent coefficients of the discrete solution of *c*_*i*_. We solve Eqs. (4) with the explicit Euler method, as solving the system is generally not particularly expensive compared to other parts of the model, hence the stringent time step requirement of the explicit Euler is not an issue here. We approximate the inverse of *M*_*II*_ using the common mass lumping technique, giving 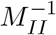 as a diagonal matrix.

To establish signaling centers, we use a simple differentiation mechanism: After each patterning step (Algorithm 1) we test if

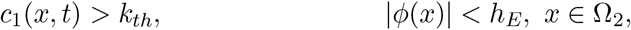

where *k*_*th*_ ∈ ℝ^+^ is the differentiation threshold and *h*_*E*_ the average mesh edge length. If the condition is satisfied, the location of the corresponding node is marked as a differentiated signaling center. In practice, multiple neighboring nodes tend to surpass the threshold simultaneously, resulting in clusters of differentiated nodes. To obtain singular signaling centers, individual differentiation centers are clustered together using a simple clique clustering, i.e., neighboring centers are clustered together if they all reside within a given (small) radius from each other. The actual signaling center position is then set as the centroid of the corresponding cluster, projected orthogonally to the interface. For simplicity, in our implementation we treat signaling centers as point sources with each localized in a single computational node.

### 2.2 Stokes flow

After obtaining growth factor distribution *c*_3_(*x*, *t*_*n*_) within tissue Ω_2_, we set a growth source term *s*(*x*, *t*_*n*_) = *c*_3_(*x*, *t*_*n*_) − *θ* with *θ* ∈ ℝ setting the zero level of the growth factor below which it acts as inducing mass sinks (i.e., apoptosis). We then solve the time-independent Stokes equation with variable viscosity and the source term (see [11]) to obtain velocity field *u*(*x*, *t*_*n*_) indicating tissue growth at time *t*_*n*_.

Given the source term *s*(*x*, *t*_*n*_), we solve for velocity *u*(*x*, *t*_*n*_) : Ω → ℝ^3^ and pressure *p*(*x*, *t*_*n*_) : Ω → ℝ with variable viscosity *μ*(*x*, *t*_*n*_) : Ω → ℝ

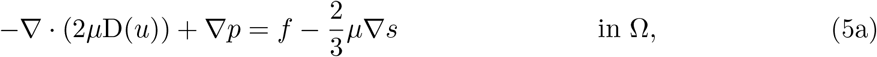

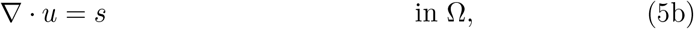

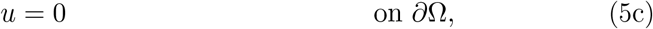

where D(*u*) is the rate-of-strain tensor defined as

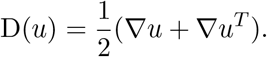

The load term *f*(*x*, *t*_*n*_) in Eq. (5a) can be used to encode the forces affecting the tissue. In this paper, we only use *f* for representing the effect of the surface tension of Γ, the purpose of which is to simulate the stiffness of the interface between two tissues, the interior and the exterior. Such stiffness could in particular apply in cases where the interface is assumed to represent an interface between two different tissue types, such as the epithelium and the mesenchyme. The use of variable viscosity *μ*(*x*, *t*_*n*_) in Eq. (5a) is to account for the changes in the adhesive properties of the tissue. For example, is has been suggested that growing tissues in some cases might have lower viscosities at growth centers [7], in which case the viscosity would be set as inversely proportional to the growth factor *c*_3_. Some possible forms of *μ* will be discussed later.

For simplicity, we set the outer domain boundary velocity to zero (Eq. (5c)). The tissue interface Γ is never supposed to come to the vicinity of *∂*Ω, hence the particular choice of boundary condition should have minimal effect on the system behavior. On the other hand, the Dirichlet boundary condition slightly reduces the computational cost of solving the system. By the divergence theorem we require for Eq. (5b) that

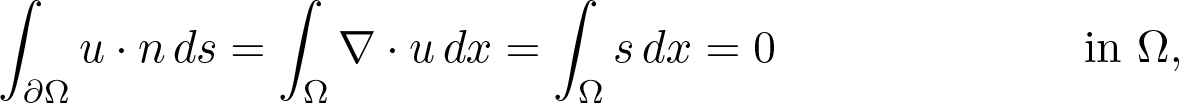

in other words, we assume that sources and sinks balance out over Ω. To obtain a unique pressure solution we impose the usual

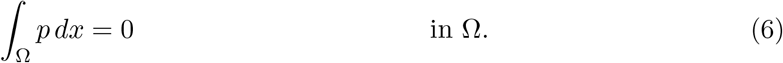

The weak form of problem (5) is: Find (*u*, *p*) ∈ *V* × *P* such that

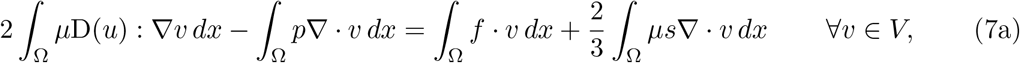

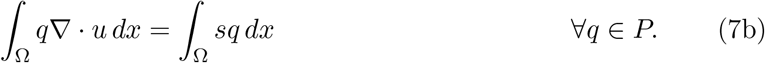

where *V* and *P* are the suitable velocity and pressure function spaces, respectively.

Solving system (7) with FEM requires care with respect to how the spaces *V* and *P* are chosen, as using the common 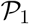 (piecewise linear) element for both spaces produces unstable pressure solution [10]. The most straightforward remedy is to use a 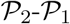 element, i.e., quadratic element for velocity and linear element for pressure, but this comes with significant computational cost. For example, using 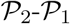 tetrahedral elements means that we will have ten computational nodes per tetrahedron (4 from vertices + 6 from edge midpoints) in contrast to the four nodes in 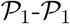 element. However, it is also possible to obtain a stable pressure solution with a reduced velocity space at some cost of numerical accuracy. A common technique is to use MINI element [3, 10]. We use a related (but distinct) stabilization through regularization suggested by Brezzi and Pitkäranta [12], which allows us to solve the system in 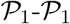 after introducing an additional stabilization term into the discrete version of system (7). The downside of this approach is that we need to determine the value of a special stabilization parameter through numerical experiments. For further discussion on stabilization, see [13].

Following [12], to obtain a stable solution, we modify Eq. (7b) for the discrete problem as

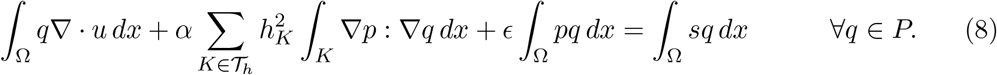

where *α* > 0 is the stabilization parameter and *h*_*K*_ the diameter of the smallest sphere containing element *K* in the set of elements 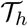 (the discretization of Ω). Eq. (8) is augmented with an additional penalty term, where *ϵ* > 0 is a small value to obtain a unique pressure solution to satisfy Eq. (6). Suitable values for parameters *α* and *ϵ* are determined later through numerical tests (see *Validation*).

Based on Eqs. (7a) and (8), assemble the following FEM matrices:

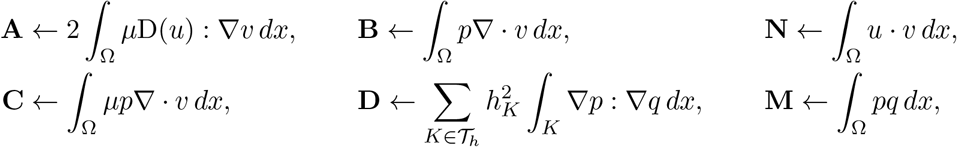

then solving the system (7a, 8) is equal to solving for the discrete solution coefficients (*u*_*h*_, *p*_*h*_) of (*u*, *p*) in a linear system

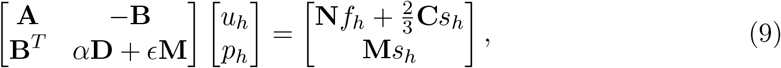

where *f*_*h*_ and *s*_*h*_ are the discrete load and source vectors, respectively. We write the right-hand side of system (9) explicitly through matrix products as it corresponds to our numerical implementation, where the matrices **N** and **M** are assembled only once and reused every time the system is solved. Further, if *μ* is constant, then **C** can be reused as well.

Presently, we only consider a load term resulting from surface tension. The normal body force acting on the interface Γ due to surface tension can be stated as

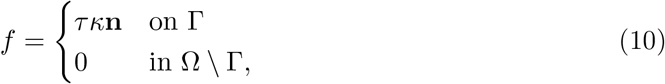

where *τ* is the surface tension coefficient, *κ* the local interface mean curvature and **n** the interface normal vector. Given the distance function *ϕ* (Eq. (1)), we have

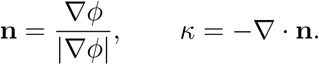

In practice, we additively distribute the values of Eq. (10) to the nodes of the edges crossed by the interface using

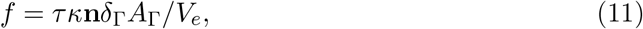

where *δ*_Γ_ is the discrete Dirac delta function with support on Γ, *A*_Γ_ the surface area of the local interface patch and *V*_*e*_ the element volume.

We solve system (9) using the iterative BiGCSTAB method [14] provided by Eigen C++ library (http://eigen.tuxfamily.org).

### 2.3 Growth

Velocity field *u*(*x*, *t*_*n*_) obtained by solving system (5) describes the flow of tissue material within Ω, and it is used to update the interface position, signaling center positions at the interface, and the morphogen concentrations due to advection.

The position of the interface Γ is updated implicitly by solving

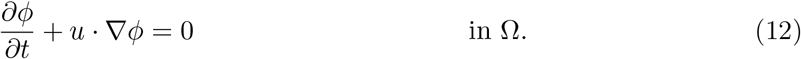

The weak form of the problem is: Find *ϕ* ∈ *V* such that

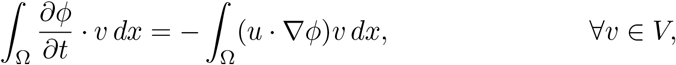

for which we use the second-order Crank-Nicolson method: Define the matrices

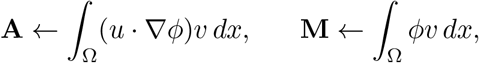

then we have an update rule

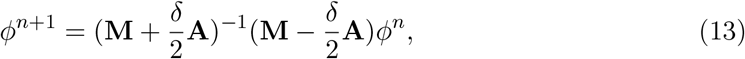

where *δ* is the time step. We approximate the velocity 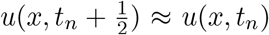 and hence assemble **A** at time *t*_*n*_, rather than at the midpoint. As in the case of the Stokes system, we solve system (13) using the BiGCSTAB method.

Solving Eq. (12) does not preserve *ϕ* as a strict distance function, and it needs to be reinitialized from time to time. Without reinitialization the interface will eventually become degenerate. A commonly cited technique for reseting the level set function is to ‘advect’ the distance values to their correct values (see, e.g., [15]). We have opted here for a straightforward computation of the minimum Euclidean distances, i.e., for a node at *x* find

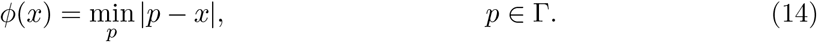

Instead of using the values obtained from Eq. (14) directly for all nodes, mass preservation can be improved by computing a weighted average of the reset (obtained form Eq. (14)) and the old pre-reset level set functions at the nodes of the edges that cross the interface. In other words, setting *ϕ*^*new*^(*x*) = *aϕ*^*old*^(*x*) + *bϕ*^*reset*^(*x*) with *a* + *b* = 1 for those *x* that belong to the edges that cross the interface. The rationale for this correction is that as the position of Γ in Eq. (14) is determined from the pre-reset level set function *ϕ*^*old*^, recomputing the distances at the interface-crossing edge nodes adds an artificial distortion to the distances, which in practice results in loss of mass. On the other hand, we found that setting the weights as *a* = 1, *b* = 0 (not recomputation) leads to accumulation of artifacts on the interface shape, especially with short advection time steps. In our implementation we use weights *a* = 3/4, *b* = 1/4.

Signaling center positions at Γ are represented by a set of points *X*(*t*) = {*X*_1_(*t*), …, *X*_*m*_(*t*)} ⊂ ℝ^3^ for *m* signaling centers. To update position *X*_*i*_ ∈ *X*, we interpolate the velocity field *u*(*x*, *t*_*n*_) at *X*_*i*_ using the barycentric coordinates: Given the enclosing element node positions *p*_*i*_ ∈ ℝ^3^, *i* ∈ 1…4, find *λ* ∈ ℝ^3^ in

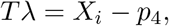

where *T* = [*p*_1_ − *p*_4_, *p*_2_ − *p*_4_, *p*_3_ − *p*_4_] ∈ ℝ^3×3^, then the interpolated velocity at *X*_*i*_ is

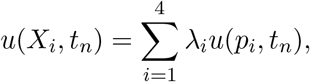

where 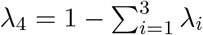. Position *X*_*i*_ is now updated as

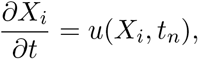

which we solve using the explicit Euler method with the same time step *δ* as the level set advection (13). Due to numerical imprecisions, signaling centers may eventually drift away from Γ, which can be remedied be re-syncing the positions *X*_*i*_ to the interface with

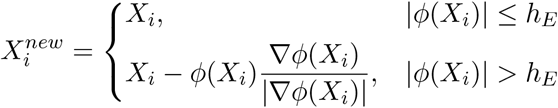

where *h*_*E*_ is the average mesh edge length. In practice, we find that the correction is only rarely needed.

The advective drift of the morphogens can be taken into account by solving

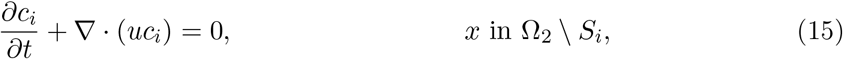

for all morphogens *i*. Note that here we have the full advective form ∇·(*uc*_*i*_) = *c*_*i*_(∇·*u*)+*u*·∇*c*_*i*_, whereas in Eq. (12) for the purpose of updating the level set function we have assumed that the divergence term has negligible effect. For updating the morphogens the full form should be used, as the concentrations fully overlap with mass sources (and sinks) within Ω_2_.

Instead of solving the morphogen advection (15), in our simulations we have opted for keeping the concentrations in place during the growth phase, and initializing the newly occupied parts of Ω_2_ to zero concentrations. This simplification is justified as we assume patterning to happen fast relative to growth and solve the patterning for long periods of time during each patterning step, minimizing the effects of zero initialization. However, in cases where growth is assumed to happen fast relative to patterning, it would be necessary to consider the advective effects of the growth to the morphogen distributions.

### 2.4 Interface reconstruction

After updating the interface position Γ, we reconstruct the implicit surface for both visualization and surface tension computation purposes. For simplicity, we assume here that the interface never coincides with the mesh edges, which in practice is true (due to floating point arithmetics) as long as the initial interface at *t*_*n*_ = 0 doesn’t coincide with the mesh edges.

The implicit surface is reconstructed by first finding all elements that cross the zero level set. Assuming that tetrahedral elements are used, each interface-crossing element has either three or four interface-crossing edges denoted by 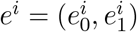, where 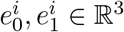 are the edge nodes. Consequently, either one or two triangles are created for the surface reconstruction (see Fig. 2). To obtain the triangle nodes, consider an edge *e*^*i*^ and the corresponding interpolated interface node

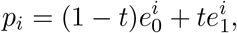

where 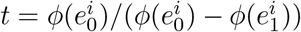. In the case of three interface-crossing edges, an interface triangle with nodes (*p*_0_, *p*_1_, *p*_2_) is created, and in the case four edges two triangles with nodes (*p*_0_, *p*_2_, *p*_3_) and (*p*_0_, *p*_1_, *p*_3_) are created.

**Fig. 2.**
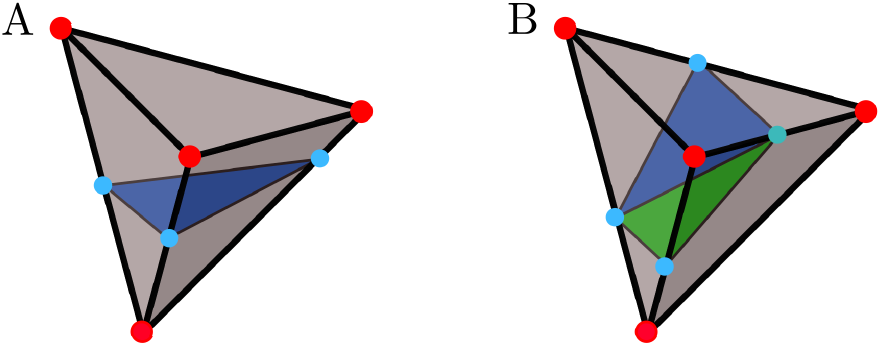
Interface reconstruction. Given an interface-crossing tetrahedral element, the interface intersects with either (A) three or (B) four element edges, with intersection points indicated by the light blue nodes. In the case of three intersection points, a single interface triangle is formed (blue). In the case of four intersection points, two interface triangles are formed (blue and green).

To obtain consistently oriented triangles for visualization, with surface normals pointing towards the exterior Ω_1_, let *p*_*ref*_ be a node of the interface-crossing element that does not coincide with the triangle nodes *p*_*i*_, and

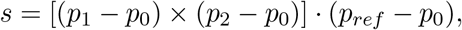

then if sgn(*s*) ≠ sgn(*ϕ*(*p*_*ref*_)), the corresponding triangle should be reversed, for example from (*p*_0_, *p*_1_, *p*_2_) to (*p*_0_, *p*_2_, *p*_1_).

## 3 Validation

To test the accuracy and correctness of the level set implementation and the Stokes solver, we simulated the effect of surface tension on an interface separating two fluids with an initial shape of a cube. The steady state solution of the free boundary problem is the sphere with volume equal to the volume of the initial shape. To further stress test the level set implementation without Stokes, we ran the Enright’s test [16] to see how well the implementation is capable of reconstructing an original shape after extreme deformation.

The effect of surface tension was simulated on a cube with the side length of 0.6 within a unit cube domain. The simulation was repeated at three levels of spatial resolution (number of nodes), 32^3^, 64^3^ and 128^3^ using the stabilized 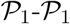 element. For reference, the simulation was further repeated at resolutions 32^3^ and 64^3^ using the stable 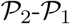 element. The simulations were run with time steps *δ* = 1/32, 1/64 and 1/128 for the three resolution levels, respectively. The total simulation time was *T* = 7/8, at which point the shape evolution was stabilized to a sphere. The main point of interest was the mass preservation over time, which was estimated using Meshlab’s Geometric Measures functionality for computing the object volume [17]. The mass loss in each case is listed in Table 1. Fig. 3A shows the simulation at three different time points at spatial resolution 128^3^ using the stabilized 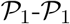 element.

**Table 1.** Loss of mass in the surface tension simulations as a function of spatial resolution.

| Resolution | Mass loss %, $\mathcal{P}_1$-$\mathcal{P}_1$ stab. | Mass loss %, $\mathcal{P}_2$-$\mathcal{P}_1$ |
| --- | --- | --- |
| $32^3$ | 8.32 | 2.89 |
| $64^3$ | 3.86 | 1.25 |
| $128^3$ | 1.67 | — |

**Fig. 3.**
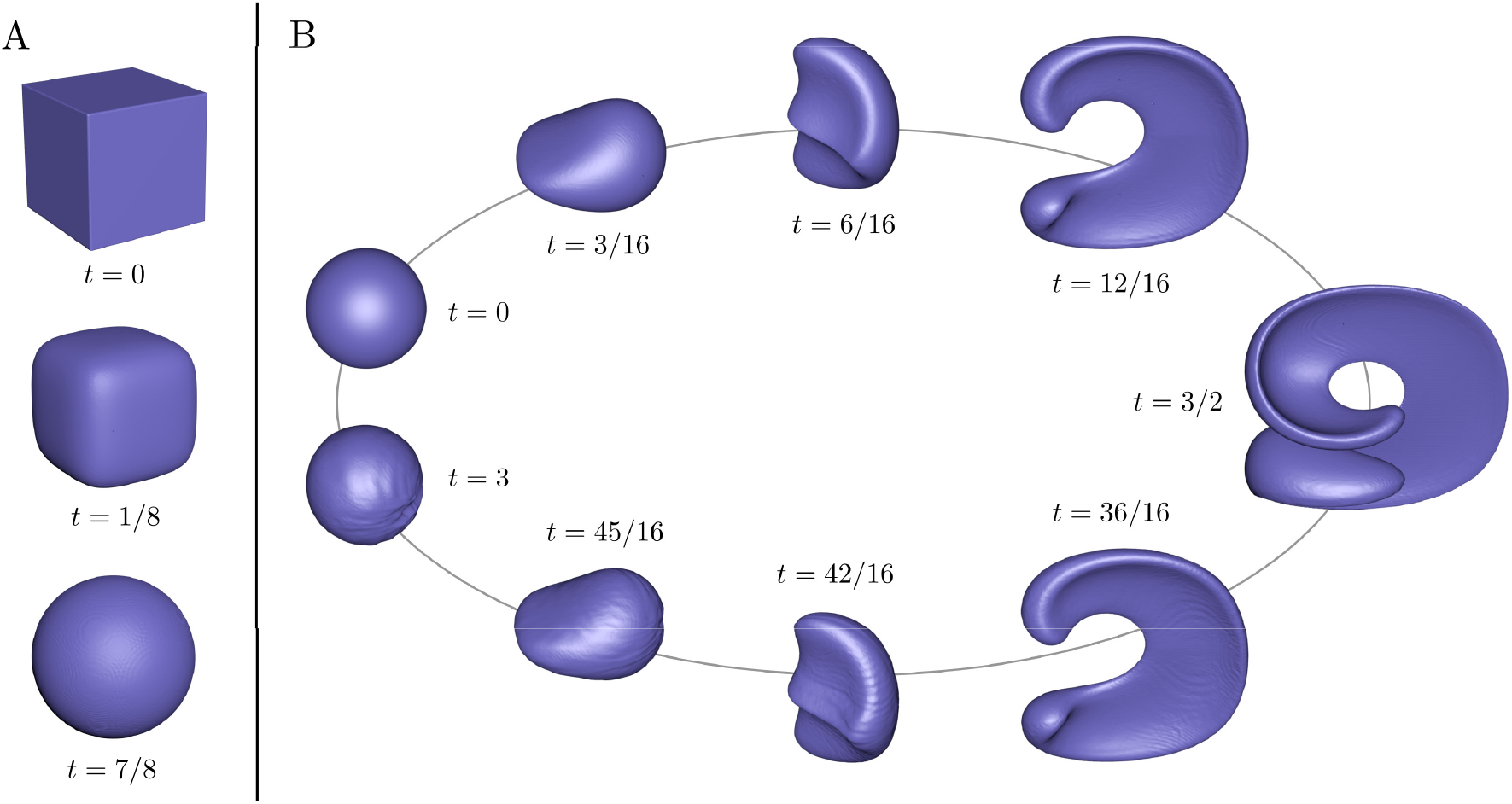
Validation testing. (A) A cube with a side length of 0.6 is driven to a sphere by imposing surface tension on the shape interface. Domain resolution 128^3^, advection time step *δ* = 1/128, surface tension coefficient *τ* = 1.0. (B) In Enright’s test, a sphere of radius 0.15 within a unit cube domain is deformed in a constant velocity field until time *t* = 3/2, after which the velocity field is reversed and the shape is returned to a sphere at time *t* = 3. Domain resolution 256^3^, advection time step *δ* = 1/64.

As might be expected, the 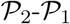 element outperforms the stabilized element by a fairly wide margin in mass preservation. However, the challenge with 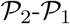 is the significantly higher computational cost: For example, in our case the simulation with 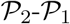 element at resolution 32^3^ took sixteen times longer compared to the stabilized 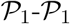 element on a laptop with i7-8750H CPU (740*s* vs. 44*s*). It is for this reason that all the growth simulations (next section) were solved using the stabilized element, as it allows for the use of higher spatial resolution, while keeping the simulation times reasonable.

In the Enright’s test a sphere with a radius of 0.15, centered at (0.35, 0.35, 0.35) within a unit cube domain, is deformed until time *T* under a velocity field suggested by LeVeque [18]

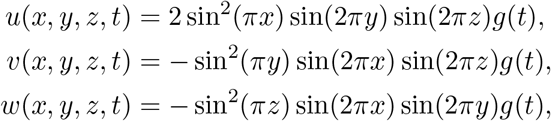

with *g*(*t*) = cos(*πt/T*). In other words, the sphere is first deformed in one direction until *t* = *T*/2, after which the time is reversed and the shape is brought back to the starting point at *t* = *T*.

We simulated the Enright’s test at three levels of spatial resolution, 64^3^, 128^3^ and 256^3^, with time steps *δ* = 1/16, 1/32 and 1/64, respectively. The total simulation time was *T* = 3. The loss of mass at each resolution level is listed in Table 2, and Fig. 3B shows the full cycle of the deformation at spatial resolution 256^3^. At the lower resolutions of 64^3^ and 128^3^ the thin film at the maximum deformation at *t* = 3/2 is partially lost, resulting in accumulation of errors and, consequently, incomplete reconstruction of the initial shape at *t* = 3. In our case the final shape was significantly deformed at the resolution 64^3^; at resolution 128^3^ the reconstruction was becoming more spherical, and at 256^3^ the shape reconstruction is fairly accurate with some ripples visible on the surface.

**Table 2.**
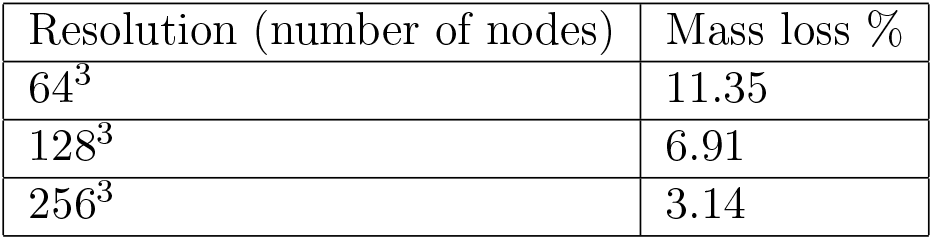
Loss of mass in the Enright’s test as a function of spatial domain resolution.

Based on the validation tests, the stabilization parameter *α* in Eq. (8) was set to 0.032 for all stabilized Stokes simulations. Using lower values of *α* generally improve the mass preservation, but also increase the spurious pressure oscillations characteristic of the unstable Stokes solution. The chosen value of *α* was found to represent a reasonable compromise in this respect. To obtain a unique pressure solution, small values in range of 10^−3^…10^−5^ were used for *ϵ* in Eq. (8). In general, the solution didn’t seem to be very sensitive to the particular choice of *ϵ*, but using a larger value of *ϵ* = 10^−3^ was found to help with obtaining good pressure solution with non-zero source terms.

## 4 Results

We first investigated the effects of surface tension and variable viscosity on the growth process starting from a small sphere. Viscosity was set as inversely proportional to the growth factor concentration: Given the maximum and minimum viscosities *μ*_*h*_ and *μ*_*l*_, respectively, we set the interior viscosity as

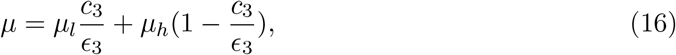

where *c*_3_ is the growth factor concentration and *ϵ*_3_ the maximum growth factor concentration at the signaling centers (see Eqs. 2). The profile is very simple in its form and used here for demonstration purposes only; see [7] for a more complex profile. The viscosity of the exterior Ω_1_ was fixed to 1.0.

Using a sphere of radius 0.15 as a starting shape and letting the patterning mechanism to establish four signaling centers in a nearly symmetrical configuration, Fig. 4A demonstrates the effect of viscosity profile (16) with *μ* ∈ [1, 100] on the growth compared to the constant viscosity *μ* = 1 in the case of zero surface tension. While the variable viscosity significantly amplifies the elongation of the growing branches, the constant viscosity also shows some branching elongation. With the addition of surface tension (Fig. 4B), constant viscosity results in near uniform ballooning of the initial initial shape, whereas the variable viscosity still demonstrates significant branching elongation.

**Fig. 4.**
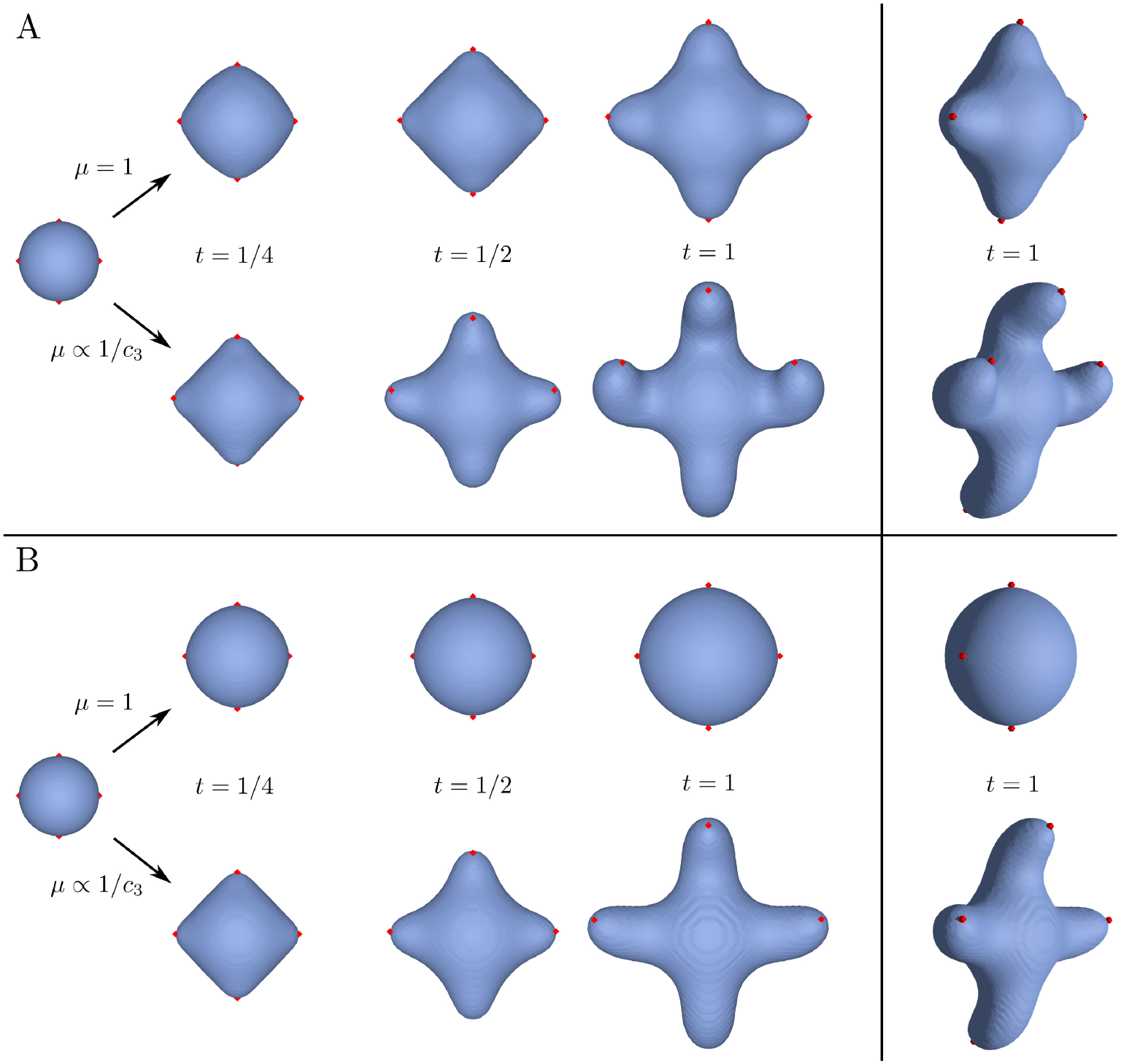
Constant vs. variable viscosity. Given signaling centers (red dots) established through the patterning mechanism (Eqs. (2)), a sphere of radius 0.15 is evolved within a unit cube domain until time *t* = 1, both (A) without surface tension *τ* = 0 and (B) with surface tension *τ* = 1.0. The upper rows in both (A) and (B) are simulated with constant viscosity *μ* = 1.0. The lower rows in both (A) and (B) have the interior viscosity set as inversely proportional to the growth factor concentration *c*_3_, with a maximum viscosity of 100.0 when *c*_3_ → 0. Domain resolution 64^3^, advection time step *δ* = 1/64. Reaction-diffusion parameters are listed in Table 3.

Next, we investigated the patterning and growth on an ellipsoid, while restricting the signaling center formation on one side of the ellipsoid. This setting is roughly motivated by the growth of mammalian molar during tooth development, where specialized signaling centers, the enamel knots, form in the epithelium near the apical surface of the dental mesenchyme and control the growth of the tooth cusps [19]. Fig. 5A demonstrates the patterning mechanism establishing the signaling centers, and Fig. 5B shows the resulting growth. The simulations were repeated with both constant and variable viscosity. Surface tension was kept at zero (no stiffness) for all simulations. As in Fig. 4, the variable viscosity is associated with amplified elongation of the growth branches, along with curving of the branches. Further, the variable viscosity causes some amount of bending of the shape base, compared to the constant viscosity case where the base simply balloons.

**Table 3.** Reaction-diffusion parameters for the growth simulations in Figs. 4–6. In Fig. 4, Eqs. (2) are solved with time step *δ* = 1/8 for 16000 iterations (total time *T* = 2) for each iteration of Algorithm 1. In Figs. 5–6, Eqs. (2) are solved with time step *δ* = 1/32 for 32000 iterations (total time *T* = 1) for each iteration of Algorithm 1.

| Parameter | Value (Fig. 4) | Value (Figs. 5–6) | Description |
| --- | --- | --- | --- |
| $\alpha_1$ | 2.0 | 2.0 | $c_1$ (activator) auto-activation |
| $\mu_1$ | $5.0 \cdot 10^{-6}$ | $5.0 \cdot 10^{-6}$ | $c_1$ diffusion rate |
| $\lambda_1$ | 0.01 | 0.01 | $c_1$ degradation rate |
| $\beta_2^1$ | 1.0 | 1.0 | $c_2$ (inhibitor) inhibition over $c_1$ |
| $\mu_2$ | $1.0 \cdot 10^{-4}$ | $7.5 \cdot 10^{-5}$ | $c_2$ diffusion rate |
| $\lambda_2$ | 0.02 | 0.02 | $c_2$ degradation rate |
| $\beta_3^1$ | 200.0 | 200.0 | $c_3$ (growth factor) inhibition over $c_1$ |
| $\mu_3$ | $1.25 \cdot 10^{-4}$ | $1.25 \cdot 10^{-4}$ | $c_3$ diffusion rate |
| $\epsilon_3$ | 40.0 | 40.0 | $c_3$ production rate |
| $\mu_3$ | 0.02 | 0.02 | $c_3$ degradation rate |
| $k_{th}$ | 7.0 | 5.0 | signaling center threshold |

**Fig. 5.**
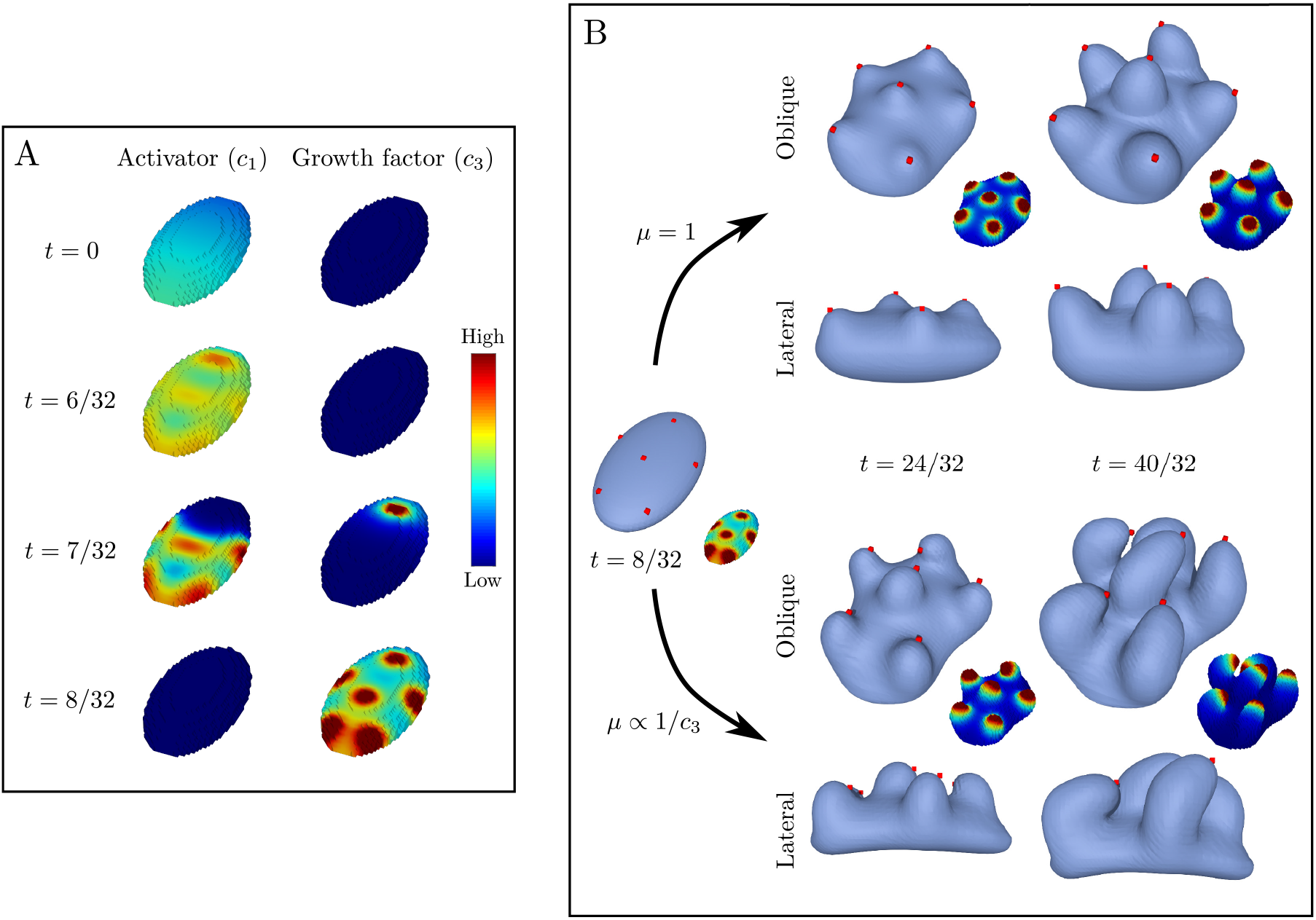
Directed growth with inverse viscosity profile. (A) The reaction-diffusion system establishes the patterning (activator peaks) that triggers the differentiation of the signaling centers that produce the growth factor. (B) Growth factor, produced by the signaling centers indicated by red dots, induces growth. On the top the viscosity is set as constant (*μ* = 1), and at the bottom inversely proportional to the growth factor concentration (*μ* ∝ 1/*c*_3_), with a maximum viscosity of 100.0 when *c*_3_ → 0. Domain resolution 64^3^, advection time step *δ* = 1/64. Reaction-diffusion parameters are listed in Table 3.

The simulations on the ellipsoid shape were repeated using a variable viscosity directly proportional to the growth factor concentration, following the observations that expression of cell adhesion molecules and components of extracellular matrix in growing tooth mesenchyme is higher near the enamel knots [20, 21]. The viscosity profile in this case had the same functional form as Eq. (16), with *μ*_*h*_ and *μ*_*l*_ switching places. The results of the simulations are shown in Fig. 6. Qualitatively, the effect of direct profile is similar to that of inverse profile in that the branching structures become elongated compared to the constant viscosity case, however, the effect is more subdued with the direct profile. Similarly, the bending of the shape base is weaker, and requires higher maximum viscosity to become visible (*μ*_*h*_ = 1000 vs. *μ*_*h*_ = 100). The direct profile was further tested with a modification where the rate of viscosity growth was set as four times faster; in this case, the bending of the base become very noticeable (Fig. 6).

**Fig. 6.**
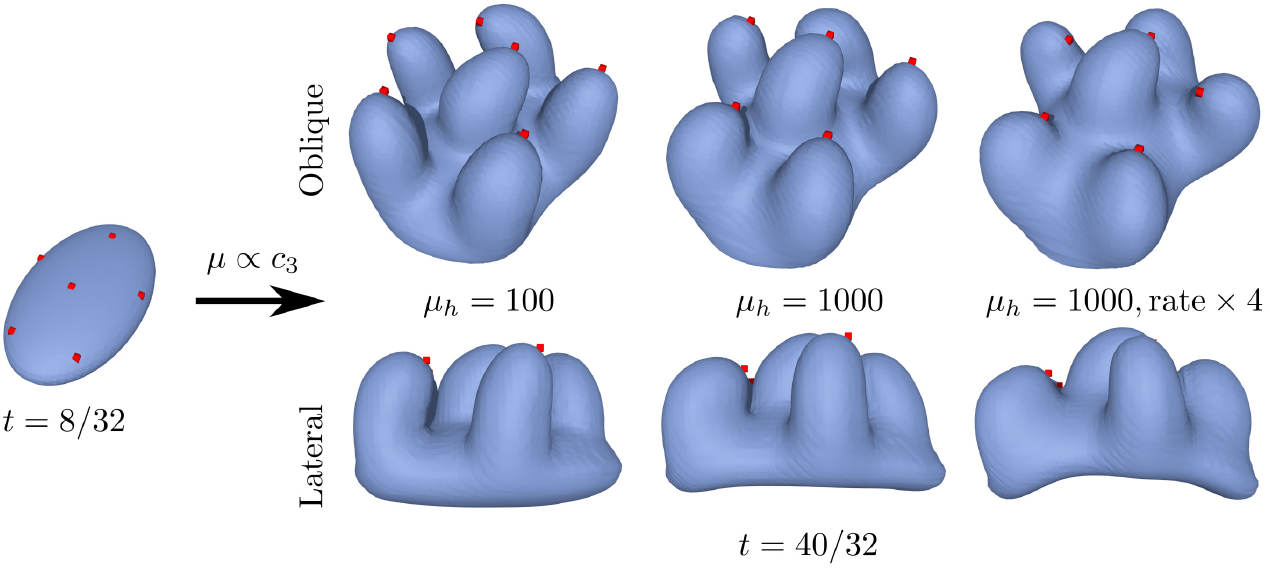
Directed growth with direct viscosity profile. Given signaling centers (red dots) established by time *t* = 8/32, growth is simulated with viscosity directly proportional to the growth factor concentration (*μ* ∝ *c*_3_). Two levels of maximum viscosity are simulated: *μ*_*h*_ = 100 and *μ*_*h*_ = 1000. An alternative viscosity profile where the viscosity grows at four times the rate relative to the growth factor is simulated for *μ*_*h*_ = 1000. Domain resolution 64^3^, advection time step *δ* = 1/64. Reaction-diffusion parameters are listed in Table 3.

All the growth simulations were performed on a laptop with i7-8750H CPU, 16GB memory and Ubuntu 18.10 operating system. The computation times varied between 20 and 190 minutes, mainly depending on the viscosity profile and the inclusion of surface tension, with high viscosity simulations combined with surface tension taking longer time solve. Solving the Stokes system (9) is the computationally most expensive part of Algorithm 1. by a wide margin, with all the other steps taking negligible time in comparison.

## 5 Discussion

We presented a computational framework that can be used to study tissue growth in cases where the tissue forms a bulk volume that can be assumed to behave mechanically as viscous fluid. We demonstrated how the Stokes flow with variable viscosity can be coupled with a dynamic patterning mechanism for establishing signaling centers to direct the tissue growth. While we have restricted our scope to simplified cases where the growth is a result of distinct patterning and growth phases, the framework is readily extendable to more complex cases where growth and patterning take place simultaneously.

Solving any Stokes problem accurately in 3D is computationally expensive with the present computer technology, and even more so when the system involves both variable viscosity and mass sources as presented here. For this reason, our focus was on the computational efficiency of the framework. Most importantly, we solve the Stokes system using a stabilized formulation, which significantly reduces both computing time and memory requirements. We track the tissue boundary using a simple level set implementation that allows us to use a static computational mesh, without the need to adaptively remesh the computational domain to fit the tissue boundary. While both of these choices imply some trade-offs in mass preservation, we demonstrated that our framework is capable of reaching reasonable accuracy even at fairly low resolution levels (see *Validation*), while being computationally sufficiently inexpensive that it can be run on ordinary desktop/laptop computers. Notably, the accuracy of mass preservation in the Enright’s test, which stress tests the level set implementation, compares favorably to even some more sophisticated level set implementations (see [22] as an example).

The growth simulations (see *Results*) demonstrated the effects of surface tension and variable viscosity on the shape formation directed by a diffusing growth factor. Surface tension had the effect of dampening the branch formation in growth, resulting from the tendency of the interface to resist curving. It could be expected that some amount of surface tension is always present in growing tissues, and in particular in cases such as developing teeth where the mesemchyme, or the bulk tissue, is largely surrounded by an epithelial layer [19]. The simplest interpretation for the variable viscosity is that cell adhesion changes throughout the tissue volume: In areas of higher cell adhesion, the flow of cells in the bulk is expected to slow down, corresponding to higher viscosity. Alternatively, forming extracellular matrix, such as collagen, may decrease tissue viscosity. However, to what extend either surface tension or variable viscosity could play role in morphogenesis is beyond the scope of this paper.

Some practical performance improvements to the numerical methods not considered here include preconditioners for the BiGCSTAB for solving the Stokes system (9). Basic diagonal/Jacobi preconditioner was tested, but was found to result in the solver failing to convergence in some instances. Currently, an unpreconditioned BiGCSTAB is used, but finding a suitable preconditioner could potentially lead to significant performance improvements [23]. The validation tests also indicated that the accuracy of mass preservation is only partly limited by the stabilized method used in solving the Stokes system, and that the accuracy could be improved by employing a more sophisticated level set implementation. Indeed, the loss of mass is a known imperfection of these interface tracking methods, and developing more accurate level set methods is an active field of research (see, e.g., [24]).

Future work will focus on applying the framework to simulating morphogenesis in real systems. Potentially suitable systems to explore include the development of molars and limbs, both of which have been targets of computational modeling in previous studies, and whose development is relatively well understood [19, 25].

## Acknowledgements

We thank Stuart Newman and Tilmann Glimm for comments and advice during the early stages of this project. Financial support was provided by the Academy of Finland and Vilho, Yrjö and Kalle Väisälä Foundation.

